# mutyper: assigning and summarizing mutation types for analyzing germline mutation spectra

**DOI:** 10.1101/2020.07.01.183392

**Authors:** William S. DeWitt

## Abstract

**Summary:** Characterization of germline mutation spectrum variation from population genomics data has shed light on the biological complexity of the mutation process, and its evolution within and between species. This analysis augments available population SNP data with estimates of local ancestral genomic context to assign mutation types and aggregate summary statistics thereof, and is increasingly common. There is a need for standardized computational tools to extract mutation spectrum information from sequencing data. Here I describe `mutyper`, a command-line utility and Python package that uses an ancestral genome estimate to assign mutation types to SNP data, compute mutation spectra for individuals, and compute sample frequency spectra resolved by mutation type for population genetic inference.

**Availability and implementation:** `mutyper` can be installed using the pip package manager and is compatible with Python 3.6+. Documentation is provided at https://harrispopgen.github.io/mutyper; source code is available at https://github.com/harrispopgen/mutyper.

## Introduction

Genomics studies of the germline mutation process seek to elucidate the mutational forces that drive genetic variation and provide the raw material for adaptive evolution. Germline mutations arise from spontaneous DNA damage caused by environmental mutagens, or errors in DNA replication. Populations and species may experience distinct histories of mutational input due to differences in environmental exposure, and heritable variation in the machinery controlling DNA replication fidelity.

Diverse mutational processes can reveal themselves in tell-tale mutation signatures left in the nucleotide sequence contexts in which they tend to be active. Population genomics has given increasing attention to nucleotide sequence context in the study of the germline mutation process (reviewed by Carlson et al.2020). SNPs can be assigned mutation context based on an estimate of the ancestral background they occurred on. It is standard to partition SNPs into *mutation types* according to ancestral and derived nucleotide states (“polarizing” the reference and alternative states), and a window of local nucleotide context in the ancestral background on which the SNP occurred. The *mutation spectrum* of an individual is defined as the relative distribution of derived state SNPs over these mutation types.

Inter- and intra-specific germline mutation spectrum variation has revealed a dynamic and evolving germline mutation process shaping modern genomic diversity. Parsing mutation spectra temporally (i.e. according to allele frequency) and spatially (i.e. in different genomic compartments) has revealed the history and present of the mutational processes, and applying such analysis to de novo mutation data may be clinically informative for rare or undiagnosed genetic diseases.

Despite many exciting findings to date, there is a lack of available software for general-purpose germline mutation type partitioning and mutation spectrum generation from population scale genomic variation data needed for larger bioinformatics pipelines or simple exploratory analysis. To address this need, I developed mutyper, an open-source command-line utility and Python package to:

- assign ancestral mutation types to SNP data in variant call format (VCF) files (Danecek et al.,2011)
- summarize mutation types at the individual level with mutation spectra
- summarize mutation types at the population level with sample frequency spectra for each mutation type
- compute distributions of ancestral *k*-mer content in genomic features specified by a BED mask file, to standardize spectra across genomic feature sets.

The literature on cancer somatic mutation signatures includes several software tools (many implemented as R packages) for clustering and dimensionality reduction that are not directly amenable to population scale germline variation data, but the package `Helmsman` (Carlson et al.,2018) enables more efficient interoperability with these tools. Complementing this integrative work, `mutyper` is a minimal, flexible, and extensible software package designed for population genomics researchers to generate the raw material needed to advance new analyses of germline mutation spectrum variation.

## Implementation

### CLI

`mutyper` is a Python package with a command-line interface (CLI) that implements the core functionality of assigning ancestral mutation types to SNPs that are input (or piped) in VCF/BCF format using the subcommand `mutyper variants`. Fast processing of VCF input is achieved with `cyvcf2` (Pedersen and Quinlan,2017), and mutation types are assigned via the INFO field for each variant, e.g. by including a key-value pair such as `mutation type=GAG>GTG.` A parameter `--k` allows the user to specify the *k*-mer context size desired (by default 3 for triplet mutation types and spectra). As in previous work, mutation types are, by default, collapsed by reverse complementation such that the ancestral state is either A or C. Alternatively, the argument `--strand file` can be used to provide a BED defining the strand orientation for nucleotide context at each site (e.g. based on direction of replication or transcription).

To polarize ancestral and derived allelic states, and define ancestral *k*-mer backgrounds, an input FASTA defining the ancestral genome estimate is required. `mutyper` uses the package `pyfaidx` (Shirley et al.,2015) for fast random access to ancestral genomic content, with minimal memory requirements. Ancestral genomes can be specified by various means. The ancestral FASTA sequence provided by the 1000 Genomes Project (1000 Genomes Project Consortium et al.,2015) was estimated from a multi-species alignment using `Ortheus` (Paten et al.,2008). In such a case, the ancestral FASTA can be passed to `mutyper` directly. Alternatively, ancestral states can be simply estimated by polarizing SNPs using an outgroup genome aligned to the reference (e.g. the chimp genome liftover to the human genome).

Below is a simple example Bash command that uses ancestral states defined in FASTA `anc.fa` to assign mutation types to SNPs in VCF file in.vcf, and redirects the augmented VCF data to `out.vcf.`

`mutyper variants anc.fa in.vcf > out.vcf`

More complex analysis can be achieved by filtering the input SNPs (i.e. with `bcftools`, Li 2011), and piping to `mutyper` instead of passing `in.vcf` as a command-line argument. The `mutyper` command-line utility is fully compatible with standard command-line pipelines for filtering SNPs or samples, masking regions, and merging/concatenating VCFs.

In addition to this core functionality, the CLI includes several other subcommands that facilitate research that aims to characterize modern mutation spectrum variation, and infer its evolutionary history. The subcommand `mutyperspectra` generates mutation spectra for each individual in input VCF/BCF data. The subcommand `mutypertargets` computes the number of ancestral genomic targets for each mutation type, optionally restricting to genomic features specified in BED format with the argument `--bed`. This facilitates standardizing mutation spectra across genomic features with varying target content. The subcommand `mutyperksfs` generates a site frequency spectrum (SFS) for each each mutation type. The SFS is a widely used population genetic summary statistic that informs both demographic history and mutation spectrum history when partitioned by mutation type (DeWitt et al.,2020).

### Python API

While the `mutyper` CLI enables one to include mutation type analyses in the context of a bioinformatics pipeline, the `mutyper` Python API enables one to perform the functions above in an interactive notebook session, or to implement custom analyses of mutation type data by interfacing with the strong ecosystem of scientific computing packages available in Python. For example, dimensionality reduction (such as principle components analysis or non-negative matrix factorization) is often used to summarize mutation spectra, and the `scikit-learn` package (Pedregosa et al.,2011) can be used in conjunction with the `mutyper` API for this purpose.

## Conclusion

The literature has seen a recent and rapid expansion of studies of germline mutation spectrum variation and evolution. This motivates the development of computational tools to make these analyses more standardized, reproducible, and accessible; `mutyper` meets this need.

## Acknowledgements

Kelley Harris and the Harris lab provided helpful comments on the software. Jedidiah Carlson and Sarah Hilton provided helpful comments on a draft manuscript.

